# Palbociclib interferes with replication origin firing in a pRb independent manner

**DOI:** 10.1101/2022.09.21.508850

**Authors:** Su-Jung Kim, Chrystelle Maric, Lina-Marie Briu, Fabien Fauchereau, Giuseppe Baldacci, Michelle Debatisse, Stéphane Koundrioukoff, Jean-Charles Cadoret

## Abstract

Over the last decade, CDK4/6 inhibitors (palbociclib, ribociclib and abemaciclib) have emerged as promising anticancer drugs. Numerous studies have demonstrated that CDK4/6 inhibitors efficiently block the pRb-E2F pathway and induce cell cycle arrest in pRb-proficient cells. Based on these studies, the inhibitors have been approved by the FDA for treatment of advanced hormonal receptor (HR) positive breast cancers in combination with hormonal therapy. However, some evidence has recently shown unexpected effects of the inhibitors, promoting needs to understand more about the mechanism of inhibitors beyond pRb. Our study demonstrates here how palbociclib impairs the origin firing in the DNA replication process in pRb-deficient cell lines. Strikingly, despite the absence of pRb, cells treated with palbociclib synthesize less DNA without any induced cell cycle arrest. Furthermore, palbociclib treatment disturbs the temporal program of DNA replication and reduces the density of replication forks. Cells treated with palbociclib show a defect in the loading of proteins of the Pre-initiation complex (Pre-IC) on chromatin, indicating a reduced initiation of DNA replication. Our findings highlight hidden effects of palbociclib on the dynamics of DNA replication and on its cytotoxic consequences on cell viability in the absence of pRb. This study provides a potential therapeutic application of palbociclib to target genomic instability towards pRb deficient patients.

**Significance Statement:** Palbociclib is a promising anticancer drug for pRb-proficient cell, particularly for hormonal receptor positive breast cancer, that induces the cell cycle arrest. But what about pRb deficient cell lines ? Our results show that Palbociclib disturb the DNA replication process inducing a replicative stress, an increase of DNA damages and leading to a significant decrease in cell viability. Palbociclib impairs the DNA synthesis reducing the number of active origins with the decrease of availability of the pre-initiation complexes. We believe that the demonstration of this effect of palbociclib on pRb-deficient cells may be a new therapeutic entry point in combination with other treatments for these types of cancer. Replicative stress can be one of weaknesses of pRb defficient cancer cells.

## Introduction

Palbociclib is a small molecule inhibitor of cyclin-dependent kinase (CDK) 4 and 6, part of the third generation of CDK inhibitors along with ribociclib and abemaciclib. They are highly selective and specific CDK4/6 inhibitors showing IC50 values in the nanomolar range *in vitro* (1). These inhibitors showed their efficacy and safety in many clinical trials (2). Based on successful outcome, the United States Food and Drug Administration (FDA) approved palbociclib as a treatment for advanced breast cancer in 2015 (https://www.fda.gov/drugs/resources-information-approved-drugs/palbociclib-ibrance)(3). Today, more than 200 clinical trials are ongoing for different types of cancer, reflecting palbociclib as a promising cancer treatment (4).

In an attempt to understand the antiproliferative molecular mechanism of palbociclib, numerous preclinical studies have investigated how palbociclib impacts downstream pathways of CDK4/6 in various cancer cell lines (5)(6). The most studied pathway is mediated by the pRb (retinoblastoma protein) that is the major substrate of CDK4/6. Indeed, pRb plays pivotal roles in the G1/S transition of the cell cycle releasing E2F family of transcription factors which in turn regulate the expression of genes involved in the progression of the cell cycle and in DNA replication (7)(8). CDK4/6 participate in this cascade as kinases phosphorylating the serine 780/795 and Serine 807/811 of pRb. Upon the treatment with palbociclib in clinical doses (around 1μM), pRb remains underphosphorylated and sequesters E2F factors, which finally induces efficient cell cycle arrest in G1(9). However, emerging evidence have shown that palbociclib has additional effects at high concentrations in a CDK4/6 or pRb independent manner (10)(11)(12)(13). To emphasize these findings, we reveal here the pRb independent actions of palbociclib on the dynamics of DNA replication at high dose.

DNA replication is a highly regulated process that ensures the duplication of genetic materials to transmit the entire genome to daughter cells. In human cells, thousands of origins are licensed through late M and early G1 phase, requiring the loading of the pre-replication complex (pre-RC) onto chromatin. Then, only a subset of licensed origins fires during S phase (14)(15). Increasing activity of S-phase related CDKs and DDK (Dbf4 dependent kinase) ensures the activation of origins by the transition from the pre-RC to the pre-initiation complex (pre-IC). Indeed, DDK, a complex of Cdc7 with Dbf4 catalytic subunit, is essential for the initiation of DNA replication (16) (17). In collaboration with S-phase CDK, DDK recruit helicase activators Cdc45 (Cell division cycle 45) and GINS (Go-Ichi-Ni-San) on chromatin leading to the formation of the active replicative helicase CMG (Cdc45-Mcm-GINS) complex (18). Defects in origins efficiency could induce replicative stress and DNA damage and threaten the accuracy and the completion of DNA replication, which may lead to genomic instability (19) (20).

In this study, we focused on two pRb deficient cancer cell lines MDA-MB-468 and NCI-H295R which are derived from either breast cancer or adrenocortical carcinoma, respectively. We demonstrate novel actions of palbociclib impairing the initiation of DNA replication. Indeed, we show that palbociclib induces an impairment of DNA synthesis during the S-phase of the cell cycle. By using different molecular approaches, we precisely decipher how and at which level palbociclib perturbs DNA replication and the stability of the genome in a pRb negative context. Thus, this study brings palbociclib into focus as a potential treatment targeting replicative stress for patients suffering from pRb deficient cancers.

## Results

### Palbociclib has impacts on cell viability in pRb deficient cell lines

To study the effect of CDK4/6 inhibitors on cellular viability, two pRb deficient cell lines, namely MDA-MB-468 and NCI-H295R, were treated with increasing doses of either palbociclib or ribociclib. Palbociclib reduced the cellular viability in a dose dependent manner while ribociclib had minimal effect on viability in both cell lines, indicating that palbociclib has additional cytotoxic effects in a pRb negative context **(Fig1.A and B)**. In order to explore the nature of the impact on cellular viability, we performed apoptosis analysis based on the activity of caspase3/7. MDA-MB-468 cells treated with 15 or 20μM of palbociclib showed a high activity of caspase3/7 whereas ribociclib had modest effect below 20μM **(Fig1.C)**. NCI-H295R cells went to apoptosis at 15μM of palbociclib and the caspase activity could not be observed at 20μM due to massive cell death **(Fig1. D)**. This finding suggests that palbociclib might have other effects in cells at relatively high doses, that do not require the presence of pRb. For all further experiments, we treated cells with 10 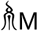 of palbociclib or ribociclib since at this concentration palbociclib reduced significantly cell viability without inducing cell death by apoptosis. Of note, cells were treated for approximately one doubling time (48h for MDA-MB-468 cells and 96h for NCI-H295R)

**Figure 1.**
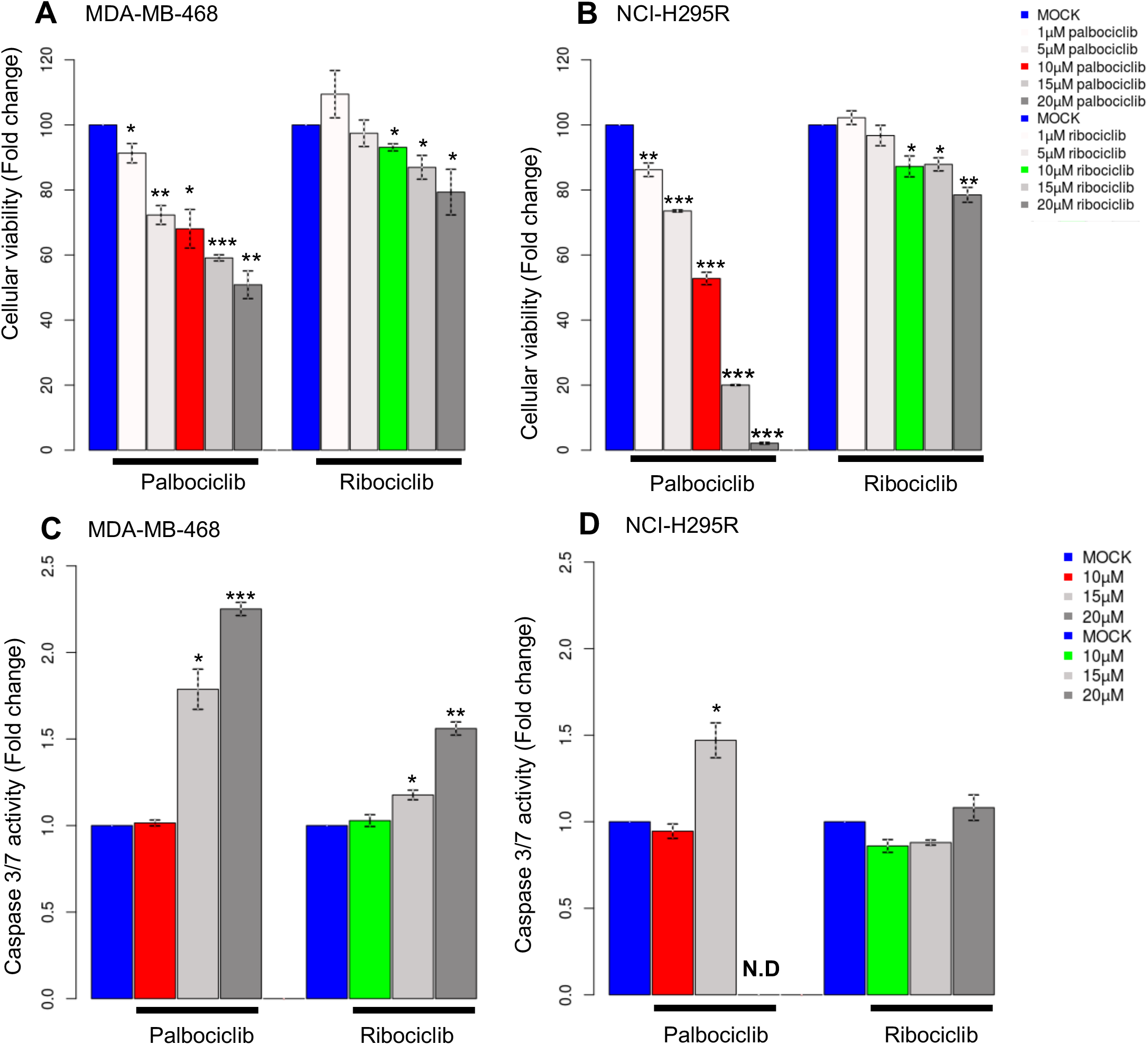
Analysis of cellular viability and apoptosis in pRb deficient cell lines. **A**. The viability of cells is measured after 48 hours of treatment with increasing doses of palbociclib or ribociclib in MDA-MB-468 cells **B**. and in NCI-H295R cells. **C**. Histograms representing the activity of caspase3/7 after treatment with palbociclib or ribociclib in MDA-MB-469 cells **D**. and in NCI-H295R cells. N.D=Non determined. Two or more independent experiments were performed for each condition (The statistical significance was tested with t-test; *P<0.05, **P<0.01, ***P<0.001).

Then, the effects of these drugs on the cell cycle were investigated. Treatment of palbociclib modestly affected the distribution of the cell cycle, with only a slight increase of the proportion of cells in G1 and G2/M phase in MDA-MB-468 and in G2/M in NCI-H295R cells **(Supplementary data 1.A and B)**. Given that the duration of treatment (one doubling time) is sufficient to observe cell cycle arrest, our results suggest that there may be an extension of the G1 or G2 with no cell cycle arrest in both cell lines.

### Palbociclib treatment impairs DNA synthesis during the S-phase of pRb deficient cells

We further analyzed more precisely the cell cycle through the assay of EdU incorporation to characterize how cells behave during the S phase of the cell cycle. This analysis allowed us to visualize the amount of newly synthesized DNA for every single cell **(Fig2.A and B)**. Comparing medians of EdU incorporation in both cell lines, we noticed that MDA-MB-468 cells incorporate more EdU than NCI-H295R cells consistently with the length of their respective doubling time **(Supplementary data 1.C)**. Strikingly, cells treated with palbociclib showed a lower intensity of EdU compared to mock treatment whereas ribociclib had minor effect on EdU incorporation **(Fig2.C)**. Palbociclib significantly reduced by 40% or 50% the level of EdU incorporation in MDA-MB-468 cells and NCI-H295R cells, respectively. Furthermore, the decrease of EdU incorporation in NCI-H295R cells treated with palbociclib was similar to the level in cells treated with 50 μM or 100μM of hydroxyurea (HU) which is considered as a mild replicative stress **(Supplementary data 2.A and B)**. These results suggest that palbociclib efficiently impairs the capacity of cells to synthesize DNA during S phase, thus implying that palbociclib may interfere with DNA replication.

Since we observed that palbociclib affects DNA synthesis, we investigated the impact of palbociclib in the presence of replicative stress and DNA damage. The activation of S-phase checkpoint was measured by the status of checkpoint Serine/threonine kinase 1 (Chk1) phosphorylation on Serine 345. We observed with the palbociclib treatment a two-fold increase in the level of phosphorylated Chk1 protein (2.41-fold in MDA-MB-468 and by 2.13-fold in NCI-H295R cells) while the total amount of Chk1 remained constant in both cell lines **(Fig 2.D and E)**. However, ribociclib showed moderate effects on the phosphorylation status of Chk1 in MDA-MB-468 cells and no effects in NCI-H295R cells. The phosphorylation level of Chk1 after treatment with palbociclib was similar to mild-replicative stress, according to the treatment with 50μM HU.

**Figure 2.**
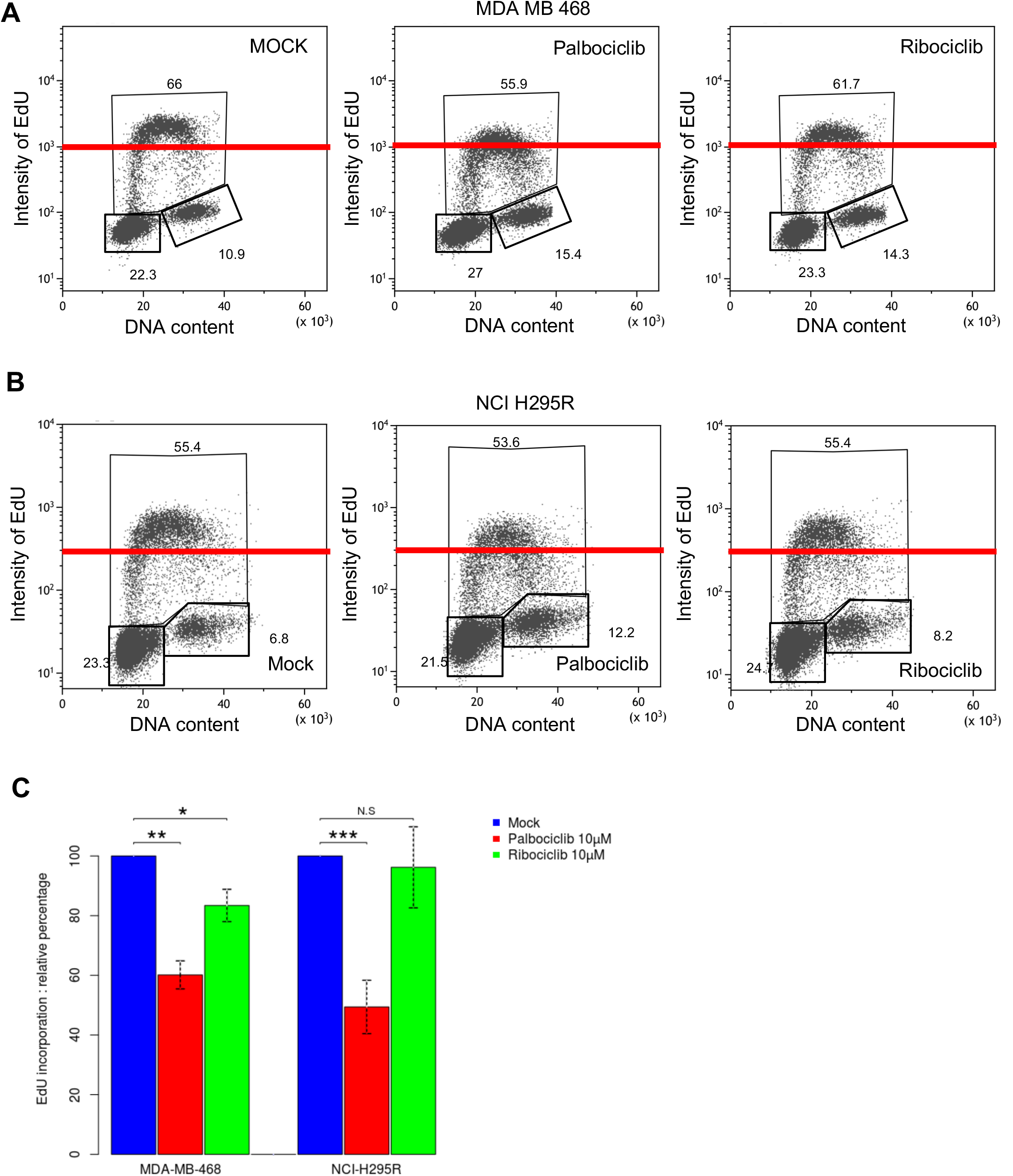
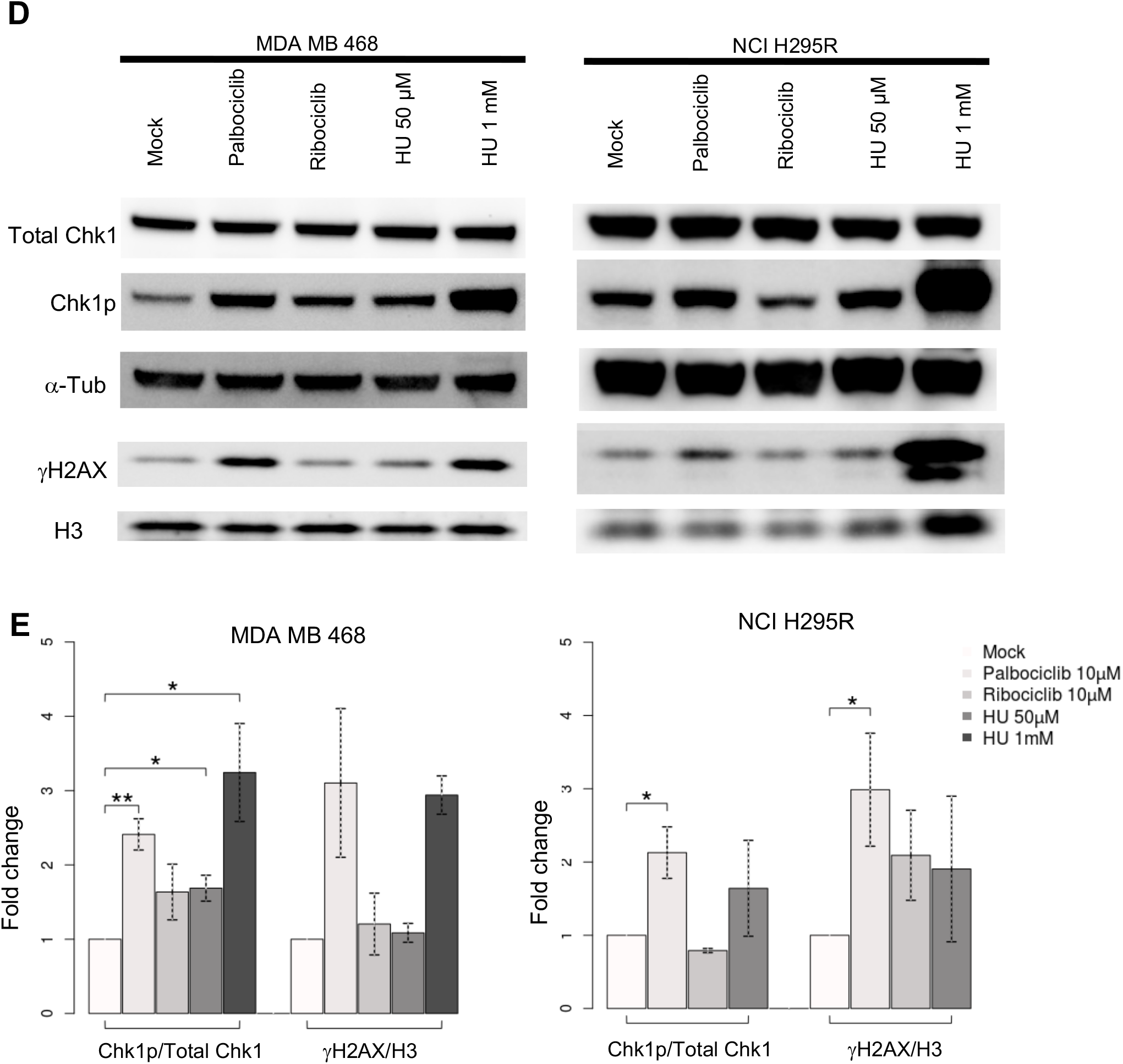
Effects of palbociclib and ribociclib on DNA synthesis during S-phase. **A**.Cell cycle profiles after EdU incorporation and drug treatment in MDA-MB-468 cells **B**. and in NCI-H295R. The red lines represent the same level of incorporated EdU for each condition. **C**. Histogram representing the relative percentage of EdU incorporated in S-phase in MDA-MB-468 and NCI-H295R cell lines. (The statistical significance was tested with t-test; *P<0.05, **P<0.01, ***P<0.001). **D**. Total protein levels of Chk1, Chk1 phosphorylated on Serine 345 residues and gH2AX were measured by Western blot after 48 hours or 96 hours of the treatments in MDA-MB-468 and NCI-H295R cells, respectively. a-Tub and histone H3 were used as loading controls. **E**. Histograms representing the relative quantity of each protein normalized to the controls. (The statistical significance was tested with t-test; *P<0.05, **P<0.01, ***P<0.001). Two or more independent experiments were performed for each analysis (C and E)

Then, we checked the induction level of DNA damage measured by γH2AX levels. We demonstrated that palbociclib triggered the accumulation of γH2AX protein whereas ribociclib had minimal effect in both cell lines. Massive DNA damages were observed in cells treated with 1mM of HU in NCI-H295R cell line. We concluded that the treatment with palbociclib drove cells to accumulate more double strand breaks in DNA. This could partially explain the modest increase of the number of cells in G2-phase that was previously observed **(Supplementary data 1.A and B)**. Cells may require a longer duration of the G2-phase to manage a complete DNA repair. Hence, cells treated with palbociclib exhibit the activation of the S-phase checkpoint and a sign of genomic instability along with interferences in DNA synthesis.

### Palbociclib reduces the number of active origins without altering replication fork progression

To decipher how palbociclib impacts on DNA replication dynamics, we performed DNA fiber analysis to measure the speed of replication forks and the activation of replication origins (21). We first examined the fork speed after a treatment with palbociclib in MDA-MB-468 cells. In control cells, the median fork speed was 2.01 kb/min **(Fig3.A and B)** and showed a slight increase (2.16kb/min), after the palbociclib treatment (although not significant, p-value=0.068). NCI-H295R cells presented a relatively low fork speed (1.18kb/min) in control condition but a statistically significant increase to 1.47kb/min after the treatment (p-value=0.028). Besides, we did not observe any difference in the replication fork asymmetry, indicating that the elongation step is not disrupted in both cell lines **(Fig3.C)**. Next, we investigated if palbociclib could impair the activation of replication origins by measuring the density of forks. After the treatment with palbociclib, the density of forks was reduced by 35% and 40% in MDA-MB-468 and NCI-H295R cells, respectively **(Fig3.D)**. It has been observed that replicative stress can induce concomitant effects on origin activation and fork speed which both influence each other (22). With regard to our data, reduced origin firing could trigger a slight acceleration of fork speed that compensate this lack of origins. Our results finally showed that palbociclib reduces the number of active origins but does not interfere with replication fork progression, which provides a mechanistic basis for the reduced EdU incorporation observed in S-phase cells by flow cytometry **(Fig. 2A and B)**.

**Figure 3.**
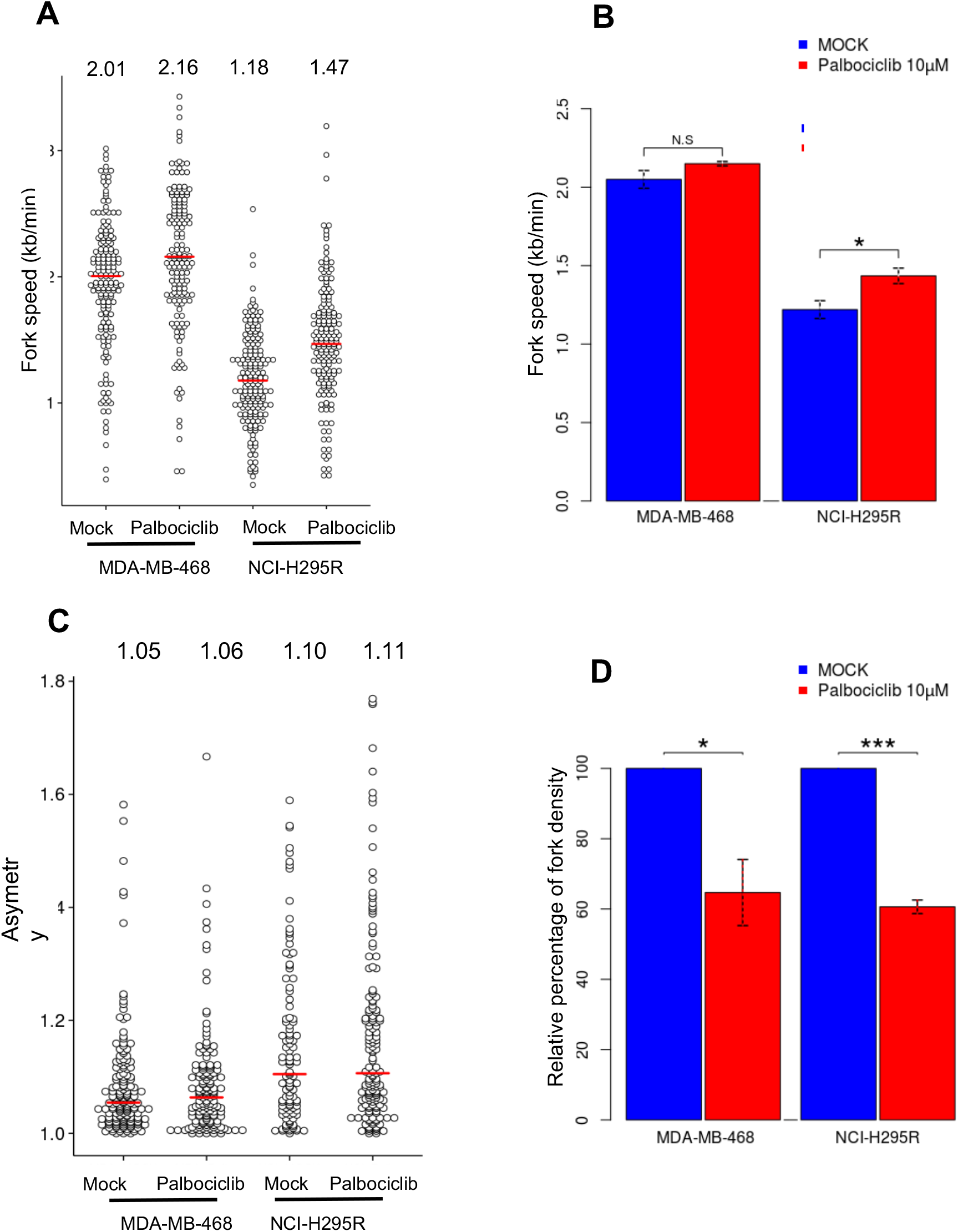
Analysis of fork speed, fork asymmetry and fork density in MDA-MB-468 and NCI-H295R cells treated with palbociclib. **A**. Dotplot indicating the fork speed of both cell lines with or without palbociclib treatment. The red lines show the median for each condition. Minimum 150 forks were scored for each condition. B. Histogram representing the fork speed mean in kb/min for each cell line. The statistical significance was tested with t-test; *P<0.05, **P<0.01, ***P<0.001) C. Fork asymmetry is measured by CldU/IdU or IdU/CldU ratios higher than 1. The red lines indicate the median of fork asymmetry. (Palbo=palbociclib treatment, Ribo=ribociclib treatment) D. Histogram representing relative percentage of fork density in cells treated with palbociclib compared to control cells treated with mock. The density of forks is calculated as the number of forks divided by the sum of the total length of the fibers. A minimum of 250kb of fibers length is counted for each condition. The statistical significance was tested as aforementioned. Two independent experiments were performed for each condition.

We observed less clusters of origins when cells are treated with palbociclib. Thus, we attempted to assess inter origin distance after treatment and found that the distance between origins is longer (169 values, 96.5kb for DMSO and 28 values, 219.7kb for Palbociclib in NCI-H295R cells). Unfortunately, we could not analyze because of lack of clusters after treatment despite of sufficient length of fibers. This observation allows us to exclude that the activation of checkpoint (Chk1p) suppresses dormant origins but palbociclib plays in the efficacity of origins.

### Palbociclib acts on the quantity of pre-initiation complex proteins

In order to better understand the molecular process, we investigated which step of the initiation of DNA replication is affected by the palbociclib treatment. We first looked at the chromatin loading of proteins constituting the Pre-Replication Complex (Pre-RC) such as Orc1, Cdt1 and Mcm2. In both MDA-MB-468 and NCI-H295R cell lines, the level of those proteins remains constant **(Fig 4.A and B)** in the different experimental conditions. We further checked the phosphorylation level of Mcm2 and the loading on chromatin of Cdc7, Dbf4 and Cdc45 implicated in the formation of pre-initiation complex (Pre-IC) **(Fig4.C)**. We found that the loaded amount of Cdc7 and Cdc45 and the level of phosphorylation of Mcm2 were decreased in the MDA-MB-468 cells after the palbociclib treatment **(Fig4.D)**. In the NCI-H295R cells as well, the same treatment efficiently decreased the loaded amount of Cdc7, Dbf4 and Cdc45 **(Fig4.E)**. These results indicate that palbociclib does not interfere with the Pre-RC loading but impairs the assembly of the pre-IC onto chromatin, resulting in reduced origins firing.

**Figure 4.**
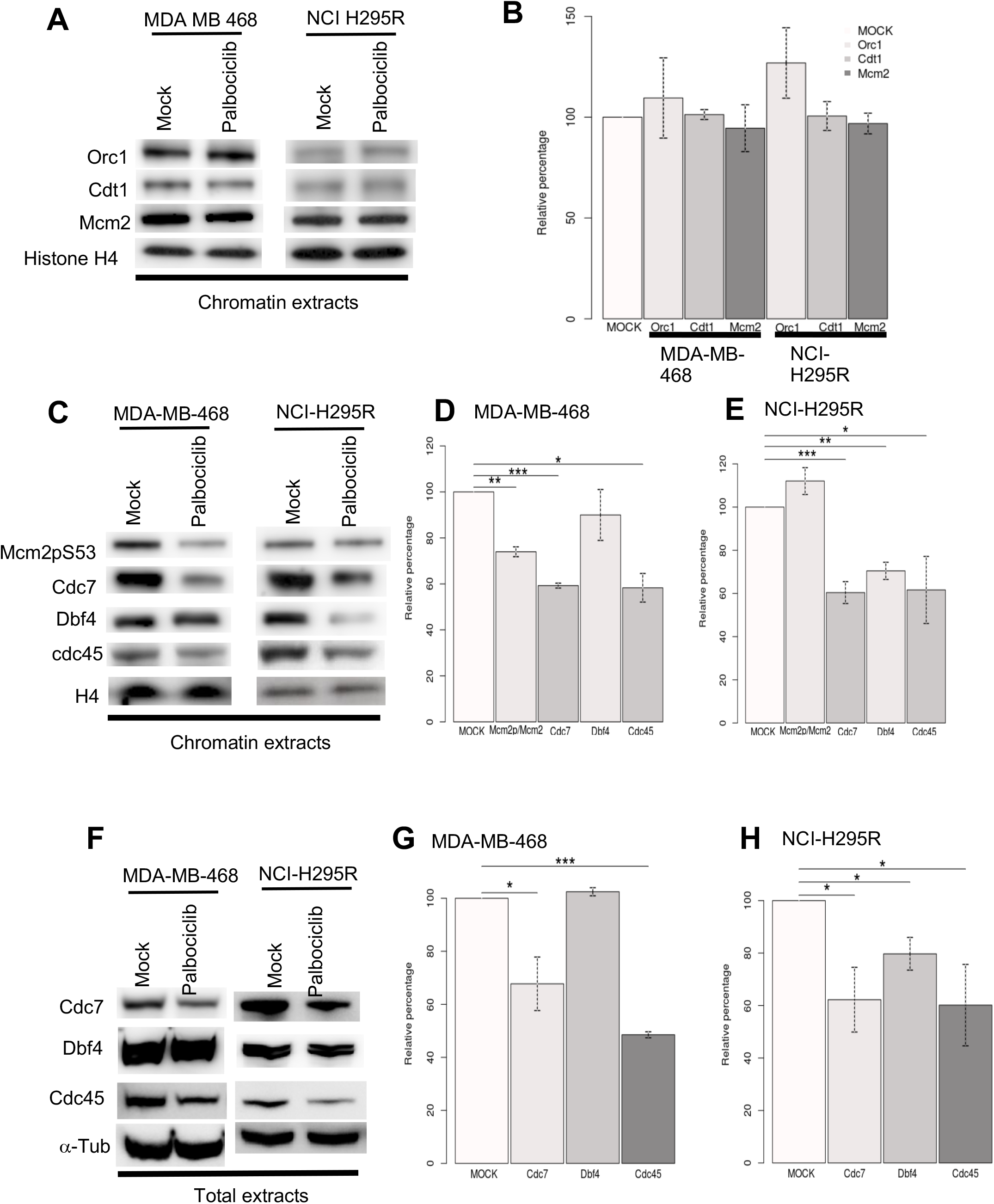
Chromatin loading of pre-ICs proteins in MDA-MB-468 and NCI-H295R cells treated with palbociclib. **A**. The fraction of proteins of the pre-RC bound to chromatin was analyzed by Western blotting. The same number of cells are used for fractionation for each condition. Histone H4 is a loading control. **B**. Same experiment as in A. for the proteins of the pre-IC. **C**. Histogram representing the relative level of proteins loaded on chromatin of the pre-RC. **D**. Histogram representing the relative level of proteins loaded on chromatin of the pre-IC in MDA-MB-468 cells. **E**. and in NCI-H295R cells. Two independent experiments were performed for each condition. (statistical significance; *P<0.05, **P<0.01, ***P<0.001). **F**. The total level of pre-initiation complex proteins was analyzed by Western blotting. The same number of cells was loaded for each condition. a-Tubulin is a loading control. **G**. Histograms representing the relative total amount of proteins in MDA-MB-468 cells. **H**. in NCI-H295R cells.

We further studied whether the decrease of chromatin bound proteins was due to a variation of the total quantity of proteins or to a defect in the pre-IC loading process. We observed that the total amounts of pre-IC proteins Cdc7 and Cdc45 proteins decreased in MDA-MB-468 cells treated with palbociclib **(Fig4.F and G)**, which is consistent with the result obtained with chromatin extracts. At the same time, less amount of Cdc7, Dbf4 and Cdc45 proteins were detected in palbociclib-treated NCI-H295R cells **(Fig4.F and H)**.

### Palbociclib regulates the expression of genes coding pre-initiation complex proteins

Then, we performed RT-qPCR to check whether palbociclib affects the mRNA level of genes coding pre-IC proteins. We detected significantly reduced levels of *Cdc7* and *Cdc45* in MDA-MB-468 cells after 24 hours and of Dbf4 and Cdc45 after 48 hours of treatment **(Fig5.A and B)**. In NCI-H295R cells, we observed that the expression levels of *Cdc7* and *Cdc45* were reduced after both 48 and 96 hours of treatment with palbociclib **(Fig5.C and D)**. Consistent with the results from analysis of proteins level **(Fig4)**, *cdc45* is repressed in a transcriptional level in two cell lines. However, we did not detect significant variation in the level of *Dbf4* whereas the protein is less present in NCI-H295R cells treated with palbociclib. It could be explained by stability of the protein due to post translational regulation (phosphorylation or ubiquitination) or binding with Cdc7 which contribute to a change in the stability of Dbf4.

By the way, we did not detect any significant variations in the level of *Orc1* and *Mcm2* in both cell lines after the treatment. Our results suggest that palbociclib regulates the expression level of specific genes particularly involved in the formation of the pre-IC in pRb-deficient cell lines. This work explains how palbociclib impairs DNA synthesis by reducing the number of activated origins without altering the replication fork progression in a pRb negative context. We revealed that the actions are mediated by transcriptional regulation of genes for the pre-IC.

### Palbociclib disturbs DNA replication process

Finally, in order to determine how palbociclib affects the DNA replication process at a genome wide scale, we performed the analysis of the temporal program of DNA replication as previously described (23). Cells organize DNA replication in space and time through the S-phase. This spatial and temporal program of replication orchestrates faithful duplication of the genetic materials during the S-phase of cell cycle. Analysis of the temporal program allows to characterize the effects of replicative stress and to visualize regions of the genome that are altered in terms of replication timing. To perform this technique, cells are labeled with a short pulse of BrdU and separated into two fractions corresponding to early and late S-phase. The ratio between these two fractions is calculated and visualized for the entire genome. Comparing the profile of control cells and cells treated with palbociclib allowed us to observe that palbociclib induced perturbations of the replication timing in several part of the genome **(Supplementary data 3.A and C)**, while the ribociclib treatment induced minimal effect **(Supplementary data 3.B and D)**.

In the MDA-MB-468 cell line, 16.82% of the genome showed an altered replication timing program after the palbociclib treatment when compared to the mock-treated control cells **(Supplementary data 3.E)**. Among the regions harboring an altered replication timing, 71.3 % displayed a delayed replication timing whereas 28.7% showed an advanced one **(Supplementary data 3.F)**. In the NCI-H295R cell line, 4.12% of the replication timing program is altered after the same treatment with 77.1% and 22.9% of delayed and advanced replicating regions, respectively. These results strengthen that palbociclib perturbs the timing of DNA replication at the genome scale **(Supplementary data 4)**.

## Discussion

Since pRb is the major phosphorylation substrate of CDK4/6, pRb loss is largely considered as a biomarker for the resistance to CDK4/6 inhibitors (4)(2). Our findings highlight the hidden effects of palbociclib using two pRb deficient cancer cell lines. In agreement with other studies, we did not detect any significant impacts of palbociclib or ribociclib around 1 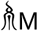 that is considered as specific inhibitory doses **(Fig1)**(5)(24). Nonetheless, palbociclib significantly reduced the cell viability at 10 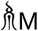, indicating that this drug has cytotoxic effects in a pRb negative context. In this study, palbociclib is used at a relatively high concentration of 10 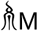. Since the effects on cellular viability are barely observed with this same concentration of ribociclib **(Fig1)**, we then suppose the actions of palbociclib are mediated by their off-targets. Recent multi-omic studies have demonstrated the different specificity of three inhibitors (ribociclib, palbociclib and abemaciclib) focusing on a large spectrum of targets for abemaciclib and also revealed that palbociclib targets other kinases than CDK4 and CDK6 at high doses (25)(13). Indeed, Willson’s group demonstrated that in live cells palbociclib shows more than 60% of occupancy to CDK2, a pivotal kinase involved in the progression of S-phase (11). Another work demonstrated that *in vitro* palbociclib also inhibits CDK2/CyclinE complex with an IC50 about 10 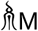 *via* a competition with the free p21 proteins, a potent cyclin-dependent kinase inhibitor (12). Their off-targets could be responsible for the effects observed in the pRB deficient cell.

In this study, we have observed that palbociclib impairs DNA replication **(Fig.2)**, the main process during the S-phase of the cell cycle. Usually Palbociclib is an efficient CDK4/6 inhibitors that induce cell cycle arrest in the G1-phase in pRb proficient cells, this cannot be the case in our study using two deficient pRB cell lines. However pRb acts not exclusively in the E2F pathway but also in chromatin remodeling by interacting with histone deacetylases (HDACs) and the ATP-dependent SWI/SNF complex (26)(27), implying that pRb deficient cells exhibit a higher intrinsic instability of the genome (28)(29)(30). Indeed our study show the unexpected observation of a decreased level in DNA synthesis after a treatment with palbociclib in pRb-deficient cell lines **(Fig2)**. The treatment in pRb deficient cells does trigger neither any cell cycle arrest nor any variation in the proportion of S-phase cells **(supplementary data 1.A and B)**, but, we noticed reduced DNA replication efficiency during the S-phase of the cell cycle **(Fig2)**. We also demonstrated the effects of palbociclib on DNA replication at the whole genome scale by studying the temporal program of DNA replication. We observed that the palbociclib disturbs the DNA replication timing, inducing significant delays in replication process in certain regions **(Supplementary data figure 3 and 4)**. We now know that p21 can interact with PCNA to inhibit DNA synthesis, particularly for the processivity of replicative DNA polymerases (*Podust VN, Podust LM, Goubin F, Ducommun B, Huebscher U (1995). “Mechanism of inhibition of proliferating cell nuclear antigen-dependent DNA synthesis by the cyclin-dependent kinase inhibitor p21”. Biochemistry. 34 (27): 8869–8875*.), and because in these cell lines p21 are free so this effect could explain the disturb of replication timing program observed in our experiments. To decipher the mechanism of action of palbociclib on the dynamics of DNA replication, we performed replication dynamic studies by the DNA fiber assay **(Figure 3)**. Our findings allowed us to eliminate the hypothesis that palbociclib affects the elongation of replication *via* p21 and PCNA interactions since no evidence has been found for fork slowing or neither fork collapse. However, the DNA combing assay indicates a reduction of the density of replicative forks by 35-40%. This decrease of the density of replication forks could be due to a defect in the initiation of origins. We first studied the loading of the pre-RC, the first complex associated with the origins of replication **(Fig. 4A and B)** and the palbociclib treatment does not affect the loading. Because the off-targets of palbociclib also affect a number of kinases, we wondered whether the activation of replication origins during the S phase *via* the pre-IC phosphorylation was impacted **(Fig. 4C, D and E)**. We detected that the total amounts of proteins constituting the pre-IC in chromatin extracts, such as Cdc7, Dbf4 and Cdc45 are reduced after treatment with palbociclib in the NCI-H295R cell line and the same is true for cdc7 and cdc45 in the MDA-MB-468 cell line (**Fig.C, D and E**) is due to the decrease of Cdc7 proteins, the DDK (Dbf4 dependent kinase) protein which activates the Pre-IC. The loading defect of Pre-IC on chromatin may also therefore be due to a hindrance of loading of Cdc7 by the presence of free p21 or/and to a decrease in the availability of Cdc45 proteins in the nucleus. To answer this question, we looked at the level of these proteins in the total extract fraction (**Fig 4F**,**G and H**). We observe for both cell lines a significant decrease in the amount of Cdc7 and Cdc45. This decrease of these protein availability is correlated with the reduction of the expression of the corresponding genes **(Figure 5A and B)**. It is well studied that a defect of origin firing could provoke an incomplete replication of the genome referred to as under replication and generate genomic instability (34). Indeed, the proteins in pre-ICs are limiting factors involved in the initiation of replication and their deregulation is tightly linked to the instability of the genome and cancer development. For instance, deregulation of Cdc7 and Cdc45 has been observed in several cancers. The depletion of Cdc7 with RNAi reduced the activation of origins in the initiation step, inducing replicative stress in the cells (35) (36). How the disturbed expression of cdc7 and cdc45 genes could occur in pRB-/-context ? It was shown that at low doses of palbociclib the cdc45 gene was inhibited *via* the Rb/E2F pathway (Knudsen ES, Hutcheson J, Vail P, Witkiewicz AK. Biological specificity of CDK4/6 inhibitors: dose response relationship, in vivo signaling, and composite response signature. Oncotarget. 2017 Jul 4;8(27):43678-43691). In addition, a non-cannonal pathway of regulation of the expression of a number of genes has been demonstrated in Rb-deficient cell lines. However, the molecular process of regulation of these genes has not yet been elucidated (The transcriptome of CDK4/6 inhibition Erik S. Knudsen, Agnieszka K. Witkiewicz Aging (Albany NY) 2017 Aug; 9(8): 1859–1860.; Knudsen ES, Hutcheson J, Vail P, Witkiewicz AK. Biological specificity of CDK4/6 inhibitors: dose response relationship, in vivo signaling, and composite response signature. Oncotarget. 2017 Jul 4;8(27):43678-43691). In our study, we can ask whether the 10-fold higher dose would not induce this non-cannonic pathway and also induce the repression of the E2F pathway despite the absence of Rb. Thus, we propose a model to explain the actions of palbociclib in inducing replication stress and DNA damages in a pRb negative context. Since the loss of pRb is associated with a higher vulnerability to replicative stress, palbociclib may potentially be used in cancer treatment with the aim of exacerbating the instability of the genome and of cancer cells becoming more vulnerable to replicative stress. From this point of view, we can consider that palbociclib may be used for pRb deficient patients combined with other therapeutic molecules that will target the replicative response pathway.

**Figure 5.**
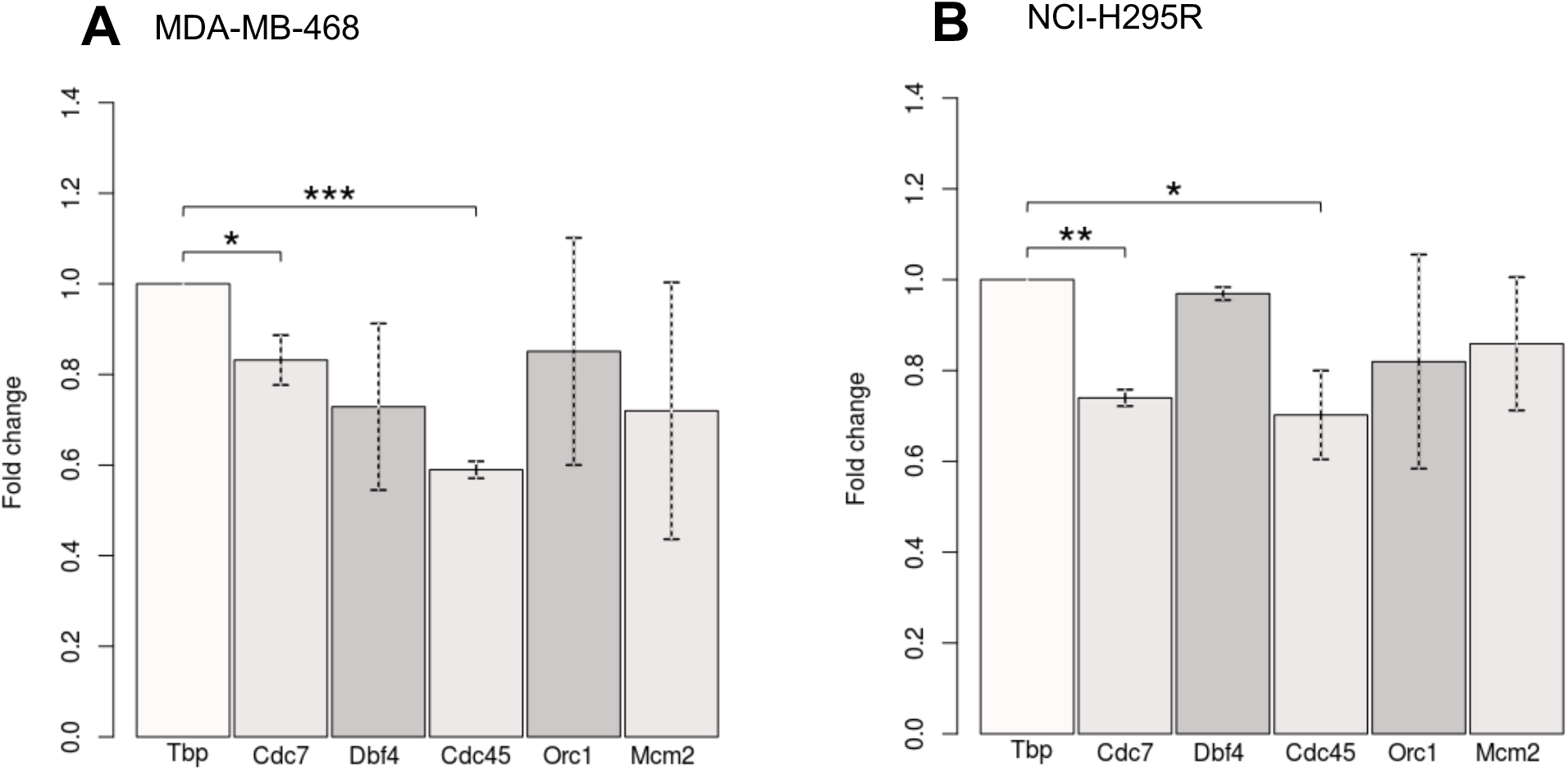
Analysis of the expression levels of Pre-IC genes. **A**. Histograms representing fold change of gene expression level for MDA-MB-468 cells treated with palbociclib for 24 hours. **B**. for NCI-H295R cells treated with palbociclib for 48 hours. The level of the Tatabox binding protein (Tbp) was considered as a control and assess at 1, from which the other genes were normalized. Two independent experiments were performed for each condition. (statistical significance; *P<0.05, **P<0.01, ***P<0.001).

## Materials and Methods

Detailed descriptions of different protocols used for this research article are provide in *SI Appendix, extended Materials and Methods*

## Acknowledgments

This project was supported by La Ligue Nationale Contre le Cancer (RS16/75-108 and RS17/75-135), the GEFLUC, the Institut National du Cancer INCa-10493, the IdEx Université de Paris ANR-18-IDEX-0001 and by the generous legacy from Ms Suzanne Larzat to our group.

## Figures and Tables

**Figure supplementary 1.**
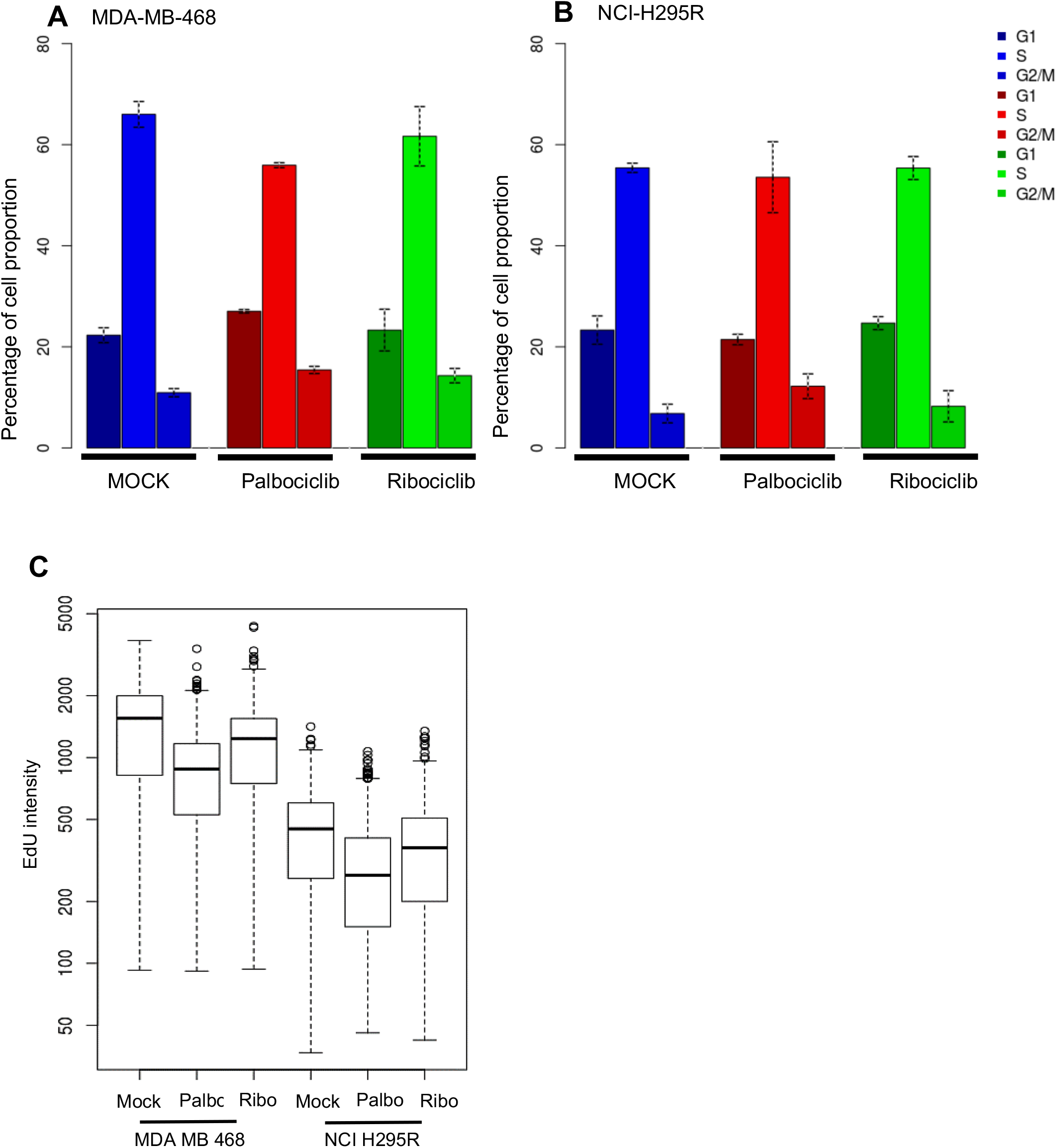
**A**. The proportion of each phase of the cell cycle is represented by percentage in MDA-MB-468 cells. **B**. and in NCI-H295R cells **C**. Boxplot representing the level of EdU incorporation in S phase in both cell lines. (Palbo=palbociclib treatment, Ribo=ribociclib treatment)

**Figure supplementary 2.**
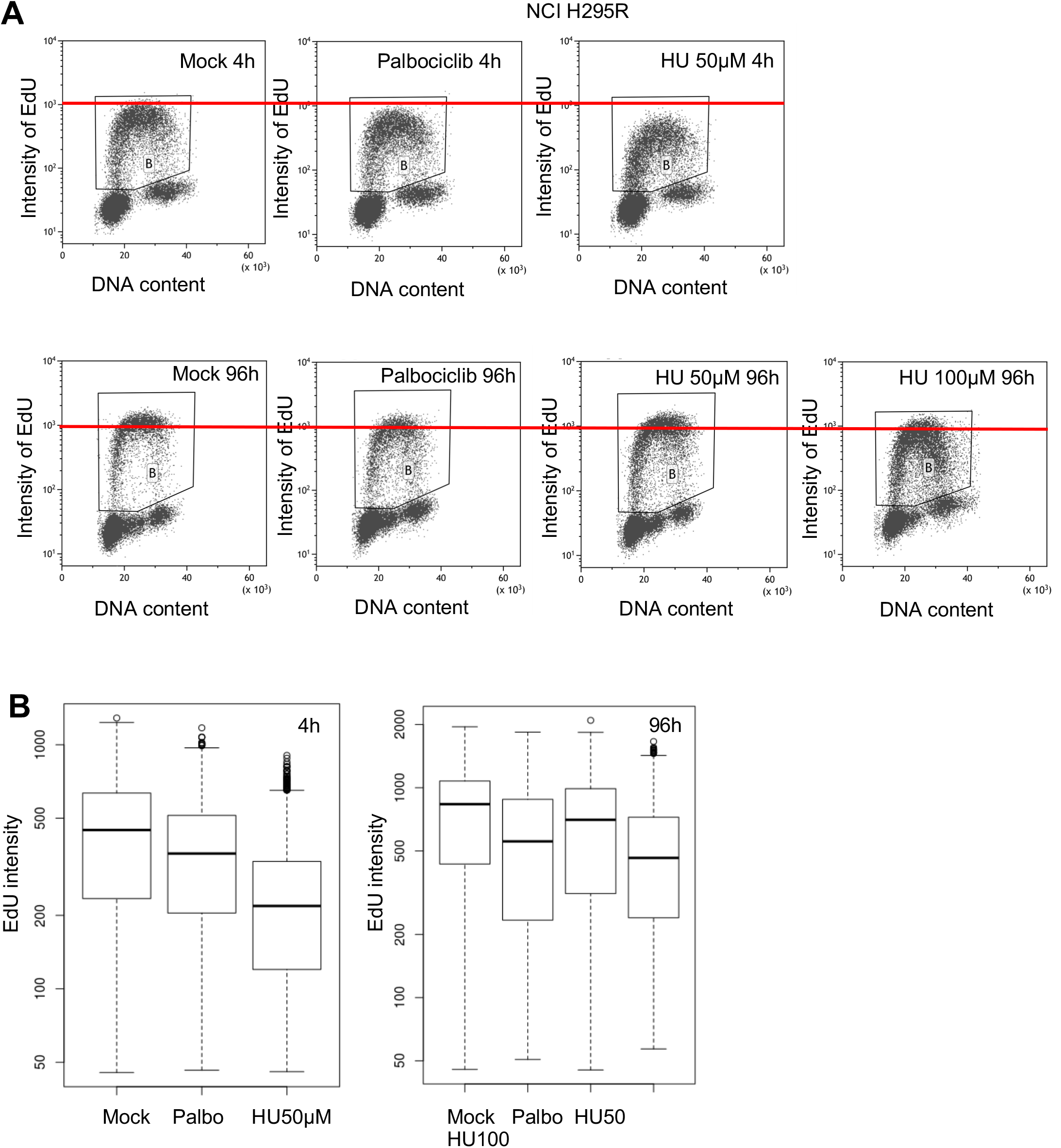
**A**. Cell cycle profiles of NCI-H295R cells after EdU incorporation and mock, palbociclib or hydroxyurea treatment for 4 or 96 hours. **B**. Boxplots representing the intensity of EdU incorporation in NCI-H295R cells in S-phase for 4 or 96 hours of treatment. (Palbo=palbociclib treatment, Ribo=ribociclib treatment, HU=Hydroxyurea)

**Figure supplementary 3.**
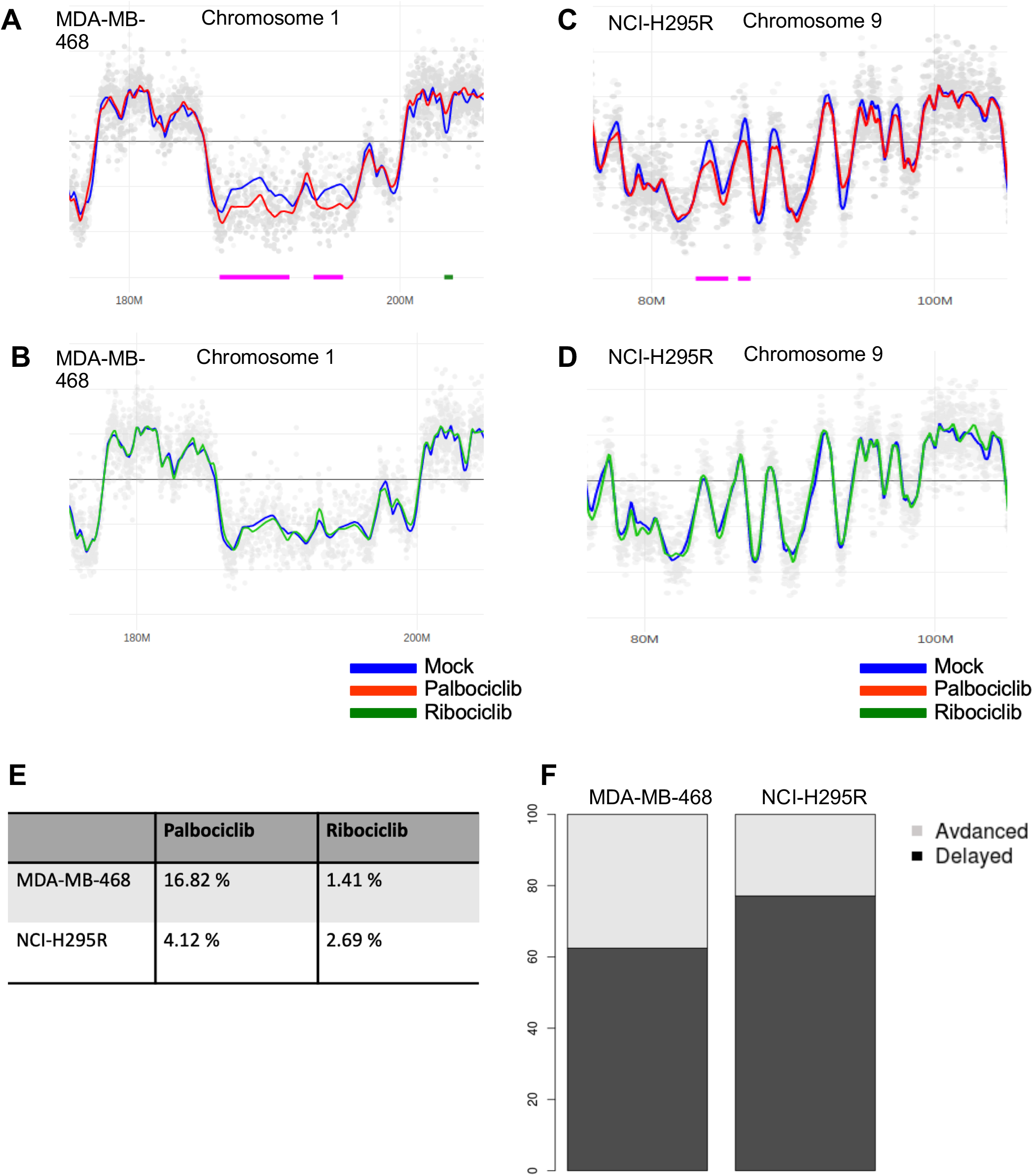
Analysis of the temporal program of DNA replication in MDA-MB-468 and NCI-H295R cells treated with palbociclib. **A and B**. Part of chromosome 1 replication timing profiles in MDA-MB-468 cells. The blue line represents control cells treated with mock, the red one for cells treated with palbociclib and the green one with ribociclib. Chromosome coordinates are indicated below each profile. **C and D**. Profile of replication timing for a part of chromosome 9 in NCI-H295R cells. **E**. Percentages of the whole genome altered after the treatment with either palbociclib or ribociclib. **F**. Stacked histograms representing the proportion of advanced or delayed replicating regions considering their length after the palbociclib treatment.

**Figure supplementary 4.**
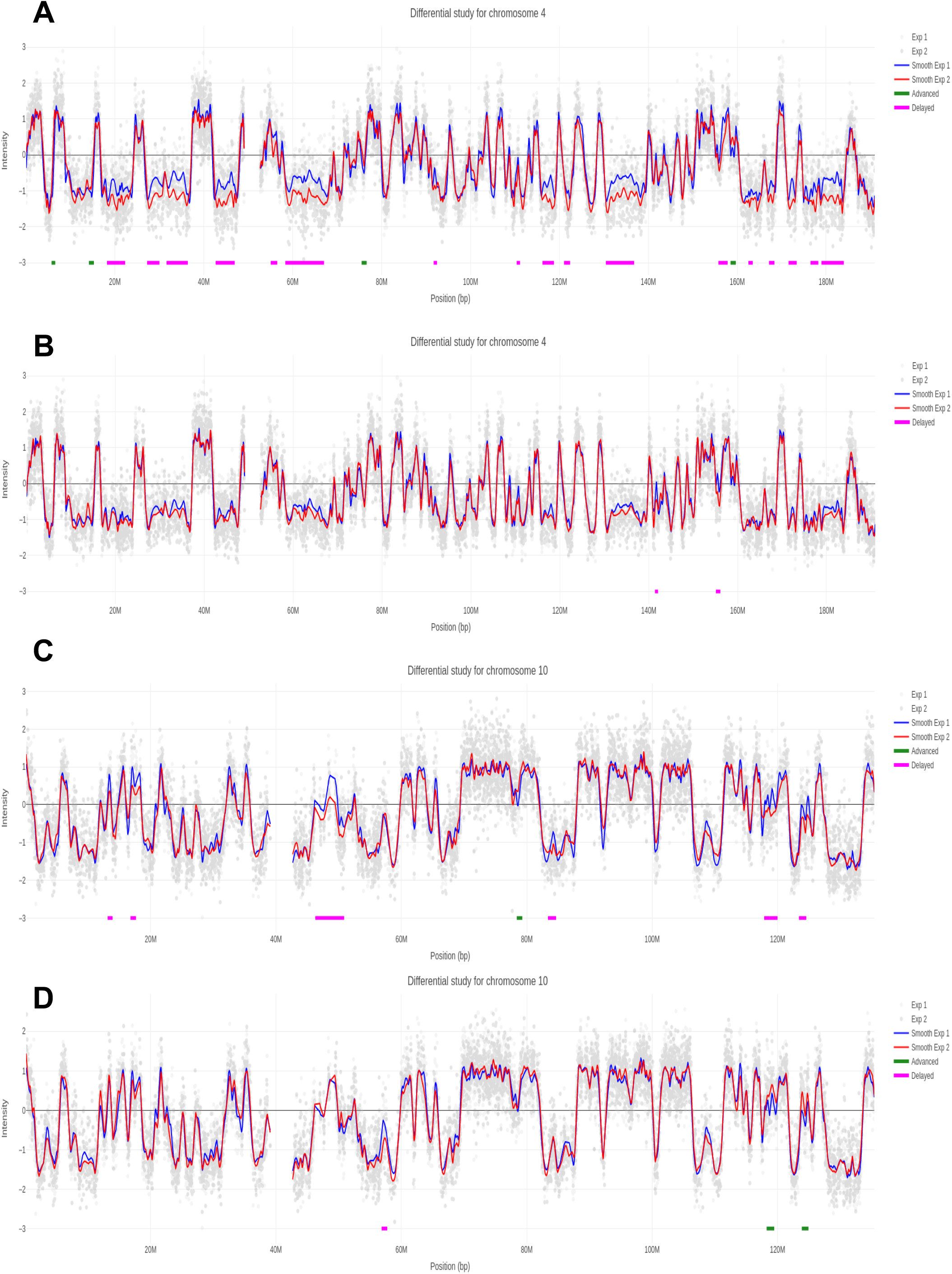
Analysis of the temporal program of DNA replication in MDA-MB-468 and NCI-H295R cells treated with either palbociclib or ribociclib. **A and B**. Chromosome 4 replication timing profiles in MDA-MB-468 cells. The blue line represents control cells treated with mock, the red one for cells treated with palbociclib in A and with ribociclib in B. Chromosome coordinates are indicated below each profile. **C and D**. Chromosome 10 replication timing profiles in NCI-H295R cells. The blue line represents control cells treated with mock, the red one for cells treated with palbociclib in A and with ribociclib in B.

**Figure supplementary 5.**
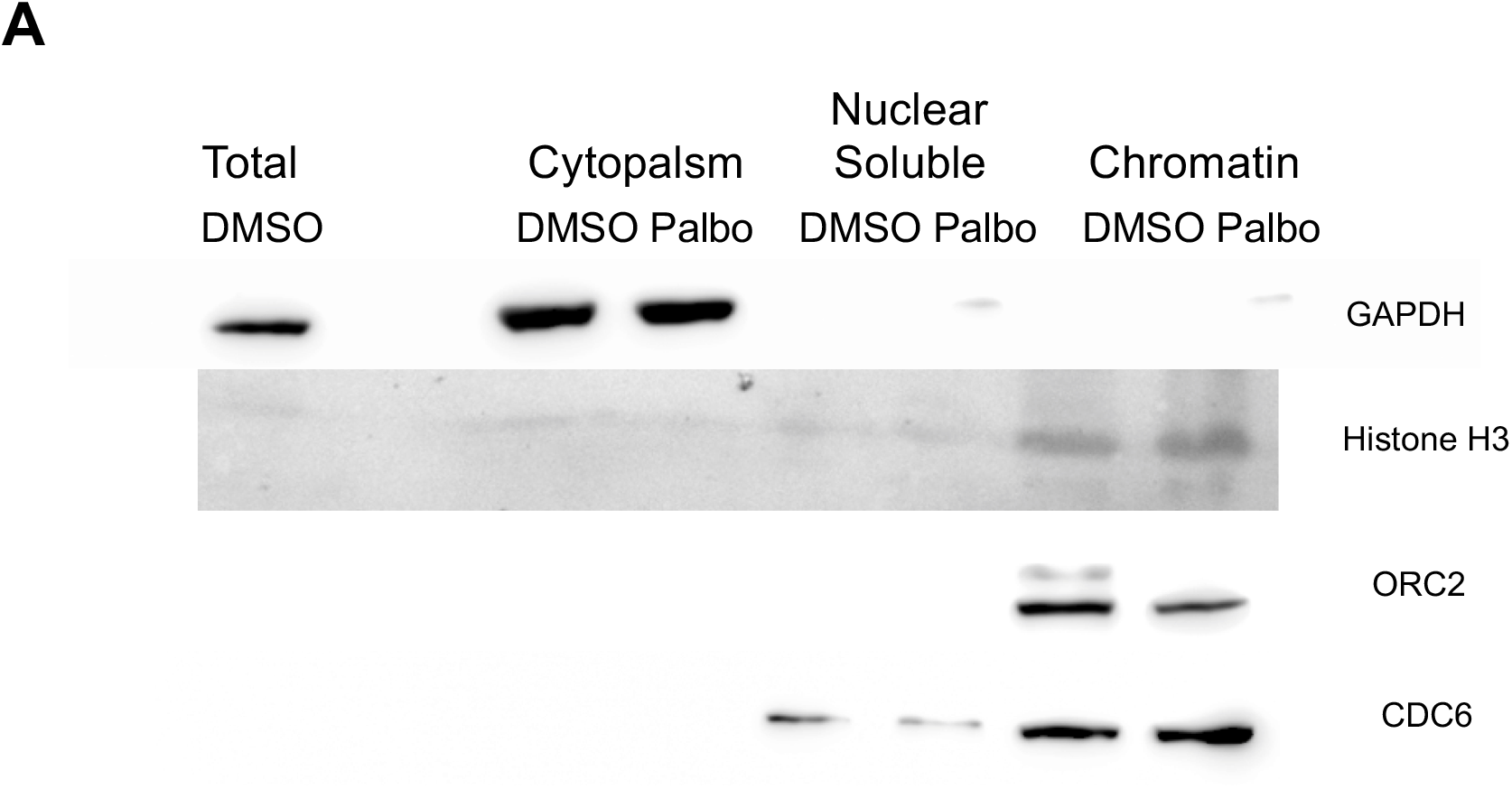
**A**. The fractions of proteins were analyzed by Western blotting. The same number of cells are used for fractionation for each condition. DMSO=96 hours of treatment with DMSO and Palbo= 96 hours of treatment in NCI-H295R cells. GAPDH and Histone H3 are used for the control of fractionation.

